# GARP complex controls Golgi physiology by stabilizing COPI machinery and Golgi v-SNAREs

**DOI:** 10.1101/2022.07.22.501184

**Authors:** Amrita Khakurel, Tetyana Kudlyk, Irina Pokrovskaya, Zinia D’Souza, Vladimir V. Lupashin

## Abstract

GARP is an evolutionary conserved heterotetrameric protein complex that is thought to tether endosome-derived vesicles and promotes their fusion in the *trans*-Golgi network. We have previously discovered the GARP’s role in maintaining Golgi glycosylation machinery. To further investigate the importance of the GARP complex for Golgi physiology, we employed Airyscan superresolution and electron microscopy, as well as the unbiased quantitative proteomic analysis of Golgi in RPE1 cells. Both *cis* and *trans*-Golgi compartments were significantly enlarged in GARP deficient cells with pronounced alterations of TGN morphology. In GARP-KO cells, proteomic analysis revealed a depletion of a subset of Golgi resident proteins, including Ca^2+^ binding proteins, glycosylation enzymes, and v-SNAREs. We validated proteomics studies and discovered that two Golgi-resident proteins SDF4 and ATP2C1, related to Golgi calcium homeostasis, as well as intra-Golgi v-SNAREs GOSR1 and BET1L, are significantly depleted in GARP-KO cells. To test if SNARE depletion is responsible for the Golgi defects in GARP deficient cells, we created and analyzed GOSR1 and BET1L KO cell lines. Since GARP-KO was more deleterious to the Golgi physiology than SNARE-KOs, we have investigated other components of intra-Golgi vesicular trafficking, particularly COPI vesicular coat and its accessory proteins. We found that COPI is partially relocalized to the ERGIC compartment in GARP-KO cells. Moreover, COPI accessory proteins GOLPH3, ARFGAP1, GBF1 were displaced from the membrane and BIG1 was relocated to endolysosomal compartment in GARP-KO cells. We propose that the dysregulation of COPI machinery along with degradation of intra-Golgi v-SNAREs and alteration of Golgi Ca^2+^ homeostasis are the major driving factors for the instability of Golgi resident proteins and glycosylation defects in GARP deficient cells.

## Introduction

The Golgi complex is a central hub in the secretory and endocytic pathway [1, 2]. It functions in trafficking, processing, and proper sorting of newly synthesized proteins in an anterograde manner while recycling Golgi-resident proteins in a retrograde manner [3, 4]. It houses multiple resident proteins, cargo receptors, sugar transporters, and glycosylation enzymes [5, 6]. The Golgi-associated retrograde protein (GARP) complex is an evolutionarily conserved multisubunit protein complex of four different subunits VPS51, VPS52, VPS53, and VPS54 [7, 8]. It is localized in the *trans*-Golgi network (TGN) and known to function in tethering the vesicles arriving from the late endosomes to the TGN [9, 10]. GARP shares its three subunits VPS51, VPS52, and VPS53 with another protein complex EARP [11]. The GARP complex is recruited to the TGN by ARL5 GTPase [12].

Mutations in GARP complex subunits have been found to cause neurodevelopmental disorders. In humans, compound heterozygous mutation of VPS51 resulted in global developmental delay, microcephaly, hypotonia, epilepsy, cortical vision impairment, pontocerebellar abnormalities, failure to thrive, liver dysfunction, lower extremity edema and dysmorphic features [13]. Protein kinase LRRK2 involved in Parkinson disease is shown to interact with VPS52 to assist in membrane fusion at the TGN thereby acting as a scaffold between GARP complex and SNAREs. Mutation in VPS52 can exacerbate Parkinson disease associated toxicity [14]. A whole exome sequencing of the genomic DNA showed compound heterozygous mutation in VPS53 leading to Progressive cerebello-cerebral atrophy type 2 (PCCA2) [15]. A missense mutation (L967Q) of VPS54 causing protein instability [16] was identified in the wobbler mouse, a model for motor neuron disease Amyotrophic lateral sclerosis [17, 18]. Moreover, a null mutation of VPS54 is embryonically lethal with high neural tube membrane blebbing phenotype [19].

Currently, it is not clear how disruption of GARP complex function is associated with severe neurodevelopmental disorders. So, to bridge this gap in knowledge, it is very important to understand the functions of GARP. GARP was initially identified to have a role sorting of Cathepsin D to lysosomes by assisting the retrieval of M6PR back to TGN [9]. Another known function of GARP is in maintenance of sphingolipids, where defects in GARP complex result in the accumulation of sphingolipids synthesis intermediates and disruption in the distribution of sterol [20]. The GARP complex is also involved in intracellular cholesterol transport via targeting NPC2 to lysosomes [21]. Auxin mediated depletion of GARP complex resulted in missorting of the flippases and remodeling of lipid composition in yeast [22]. We have recently shown that a knock-out (KO) of the GARP complex subunits affects core Golgi function of *N*- and *O*-glycosylation as a result of reduction in total level of Golgi enzymes responsible for Golgi glycosylation [23].

In this study, we continue the investigation of the role of GARP in Golgi homeostasis by employing microscopy approaches and quantitative proteomics of isolated Golgi membranes and characterizing effects of GARP depletion on Golgi Ca^2+^ binding proteins, SNAREs and COPI vesicle budding machinery.

## Methods

### Cell culture

hTERT RPE1 wild-type and GARP mutants were described previously [23]. Cells were cultured in DMEM containing Nutrient mixture F-12 (Corning) supplemented with 10% fetal bovine serum (FBS) (Thermo Fisher) and incubated in a 37°C incubator with 5% CO2 and 90% humidity.

### Cell fractionation

hTERT-RPE1 VPS54KO, VPS54KO R, VPS53KO, and VPS53KO R cells were plated in two 15 cm dishes in DMEM/F12 medium containing 10% FBS and grown to 100% confluence. Cells were washed twice with 10 ml of PBS and once with hypertonic 0.25 M sucrose in PBS. After the complete removal of PBS, the dishes were placed on ice and 1 ml of freshly prepared hypotonic ice-cold lysis buffer (20 mM HEPES pH 7.4, 1 mM PMSF, 5 µl/ml of HALT protease inhibitor) was added. The cells were collected with cell scrapper and moved to 1.5 ml microcentrifuge tube. Cells were lysed by passing through 26G syringe for 20 passages. The efficiency of lysis was examined by a phase contrast microscope equipped with 10x objective. Cell lysates were centrifuged at 3,000 g for 3 min at 4°C to pellet unlysed cells and nucleus. The S3 supernatant was transferred to Beckman 1.5 ml ultracentrifuge tube and subjected to centrifugation at 30,000 g for 30 min at 4°C. The supernatant (S30) was carefully removed without disturbing the pellet. The pellet (P30) was resuspended in 350 µl of 50% Nycodenz in the lysis buffer and transferred to 2.2 ml Beckman centrifuge tube on top of 100 µ l of 60% Nycodenz. The sample was then overlaid with 35%, 30%, 25% and 20% Nycodenz (400 µl of each), and finally with 100 µl of lysis buffer. The gradient was centrifuged at 214000 g for 4 h at 4°C. 10 fractions (200 µl each) were collected from the top, mixed with 2x SDS sample buffer, heated at 70°C for 10 min and analyzed by western blot.

To prepare Golgi-enriched membranes for Mass spectrometric analysis, membranes floated in Nycodenz gradient (top 500 µl fraction) were collected, diluted with 1 ml of 20 mM HEPES with 150 mM NaCl and pelleted at 120000 g for 1 h at 4°C. The supernatant was removed, and the Golgi membrane pellets were submitted for label-free mass spectrometry analysis.

### Mass-spectrometry and Data analysis

Proteins were reduced, alkylated, and purified by chloroform/methanol extraction prior to digestion with sequencing grade modified porcine trypsin (Promega). Tryptic peptides were then separated by reverse phase XSelect CSH C18 2.5 um resin (Waters) on an in-line 150 × 0.075 mm column using an UltiMate 3000 RSLCnano system (Thermo). Peptides were eluted using a 60 min gradient from 98:2 to 65:35 buffer A:B ratio. Eluted peptides were ionized by electrospray (2.2 kV) followed by mass spectrometric analysis on an Orbitrap Exploris 480 mass spectrometer (Thermo). To assemble a chromatogram library, six gas-phase fractions were acquired on the Orbitrap Exploris with 4 m/z DIA spectra (4 m/z precursor isolation windows at 30,000 resolution, normalized AGC target 100%, maximum inject time 66 ms) using a staggered window pattern from narrow mass ranges using optimized window placements. Precursor spectra were acquired after each DIA duty cycle, spanning the m/z range of the gas-phase fraction (i.e., 496-602 m/z, 60,000 resolution, normalized AGC target 100%, maximum injection time 50 ms). For wide-window acquisitions, the Orbitrap Exploris was configured to acquire a precursor scan (385-1015 m/z, 60,000 resolution, normalized AGC target 100%, maximum injection time 50 ms) followed by 50x 12 m/z DIA spectra (12 m/z precursor isolation windows at 15,000 resolution, normalized AGC target 100%, maximum injection time 33 ms) using a staggered window pattern with optimized window placements. Precursor spectra were acquired after each DIA duty cycle.

Buffer A = 0.1% formic acid, 0.5% acetonitrile

Buffer B = 0.1% formic acid, 99.9% acetonitrile

Following data acquisition, data were searched using an empirically corrected library and a quantitative analysis was performed to obtain a comprehensive proteomic profile. Proteins were identified and quantified using EncyclopeDIA [24] and visualized with Scaffold DIA using 1% false discovery thresholds at both the protein and peptide level. Protein exclusive intensity values were assessed for quality using an in-house ProteiNorm app, a tool for systematic evaluation of normalization methods, imputation of missing values and comparisons of multiple differential abundance methods [25]. Normalization methods evaluated included log2 normalization (Log2), median normalization (Median), mean normalization (Mean), variance stabilizing normalization (VSN) [26], quantile normalization (Quantile) [27], cyclic loess normalization (Cyclic Loess) [28], global robust linear regression normalization (RLR) [29], and global intensity normalization (Global Intensity) [29]. The individual performance of each method was evaluated by comparing of the following metrices: total intensity, pooled intragroup coefficient of variation (PCV), pooled intragroup median absolute deviation (PMAD), pooled intragroup estimate of variance (PEV), intragroup correlation, sample correlation heatmap (Pearson), and log2-ratio distributions. The normalized data were used to perform statistical analysis using linear models for microarray data (limma) with empirical Bayes (eBayes) smoothing to the standard errors [28]. Proteins with an FDR adjusted p-value < 0.05 and a fold change > 2 were considered significant.

### Western blot analysis

Total cell lysates were prepared as described before [23]. Briefly, cells grown on tissue culture dishes were washed twice with PBS and lysed in 2% SDS preheated at 70°C. Cell lysates were collected and briefly sonicated to break chromosomal DNA. Protein concentration was measured using BCA protein assay (Pierce). 6× SDS sample buffer containing beta-mercaptoethanol was added and samples were heated at 70°C for 10 min. Samples (10–30 µg of protein) were loaded onto Bio-Rad (4–15%) or Genescript (8–16%) gradient gels. Proteins were transferred to nitrocellulose membranes using the Thermo Scientific Pierce G2 Fast Blotter. Membranes were washed in PBS, blocked in Odyssey blocking buffer (LI-COR) for 20 min, and incubated with primary antibodies for 1 h at room temperature (RT) or overnight at 4°C. Membranes were washed with PBS and incubated with secondary fluorescently tagged antibodies diluted in Odyssey blocking buffer for 60 min. All the primary and secondary antibodies used in the study are listed in Table 1. Blots were then washed and imaged using the Odyssey Imaging System. Images were processed using the LI-COR Image Studio software.

**Table 1:**
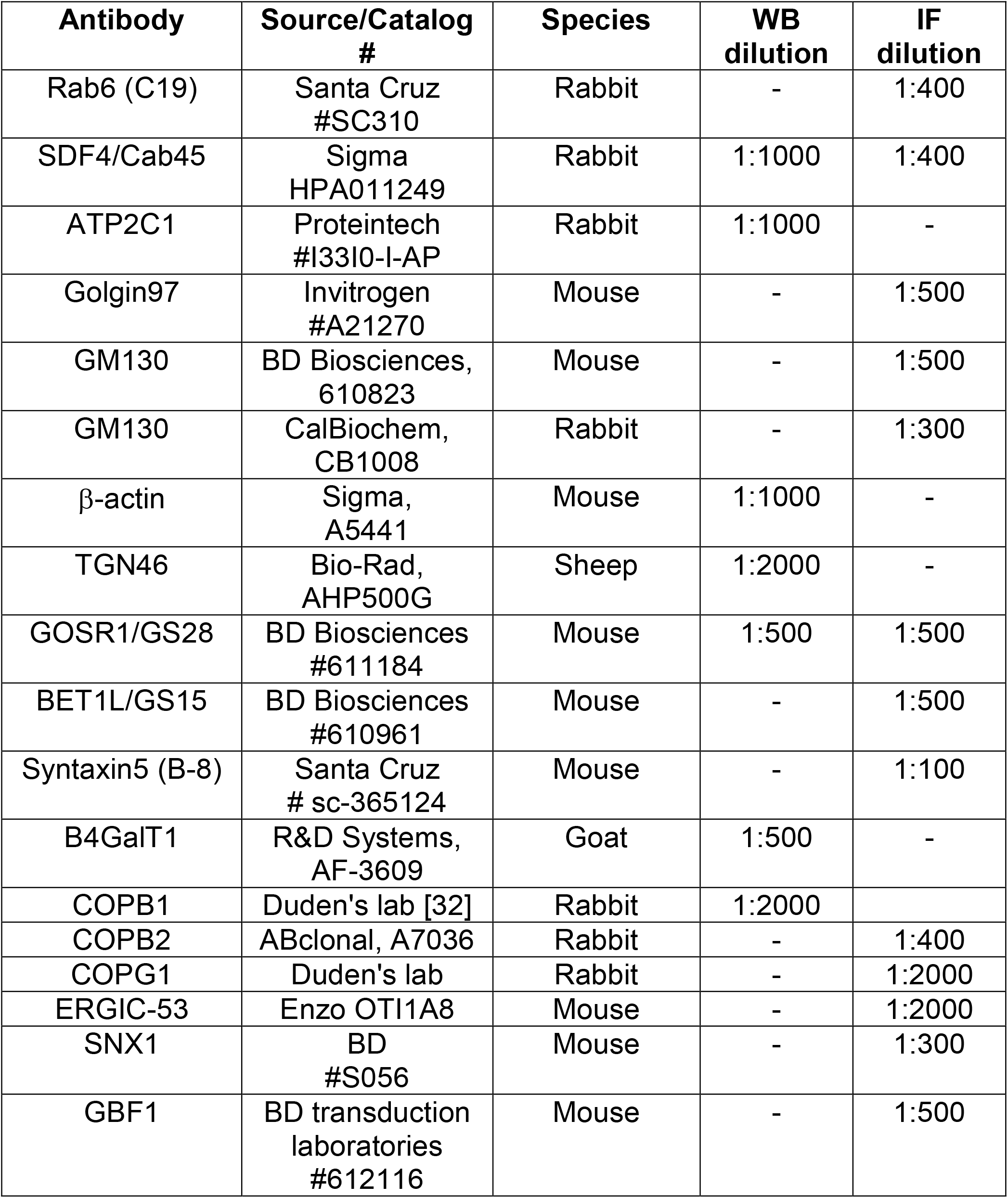
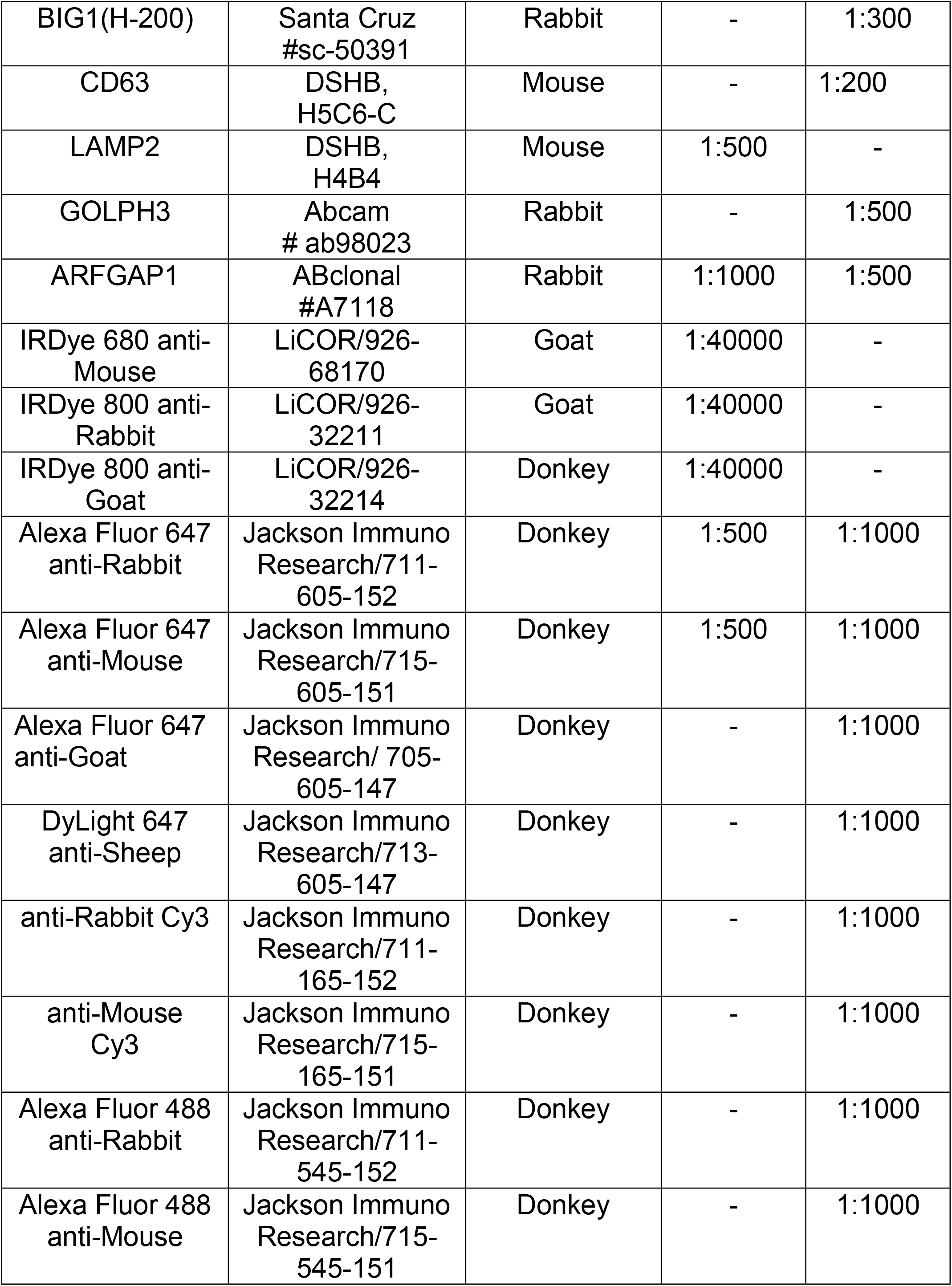
List of antibodies used

| **Antibody** | **Source/Catalog #** | **Species** | **WB dilution** | **IF dilution** |
| --- | --- | --- | --- | --- |
| Rab6 (C19) | Santa Cruz #SC310 | Rabbit | - | 1:400 |
| SDF4/Cab45 | Sigma HPA011249 | Rabbit | 1:1000 | 1:400 |
| ATP2C1 | Proteintech #13310-I-AP | Rabbit | 1:1000 | - |
| Golgin97 | Invitrogen #A21270 | Mouse | - | 1:500 |
| GM130 | BD Biosciences, 610823 | Mouse | - | 1:500 |
| GM130 | CalBiochem, CB1008 | Rabbit | - | 1:300 |
| β-actin | Sigma, A5441 | Mouse | 1:1000 | - |
| TGN46 | Bio-Rad, AHP500G | Sheep | 1:2000 | - |
| GOSR1/GS28 | BD Biosciences #611184 | Mouse | 1:500 | 1:500 |
| BET1L/GS15 | BD Biosciences #610961 | Mouse | - | 1:500 |
| Syntaxin5 (B-8) | Santa Cruz # sc-365124 | Mouse | - | 1:100 |
| B4GalT1 | R&D Systems, AF-3609 | Goat | 1:500 | - |
| COPB1 | Duden's lab [32] | Rabbit | 1:2000 |  |
| COPB2 | ABclonal, A7036 | Rabbit | - | 1:400 |
| COPG1 | Duden's lab | Rabbit | - | 1:2000 |
| ERGIC-53 | Enzo OTI1A8 | Mouse | - | 1:2000 |
| SNX1 | BD #S056 | Mouse | - | 1:300 |
| GBF1 | BD transduction laboratories #612116 | Mouse | - | 1:500 |
| BIG1(H-200) | Santa Cruz<br>#sc-50391 | Rabbit | - | 1:300 |
| CD63 | DSHB,<br>H5C6-C | Mouse | - | 1:200 |
| LAMP2 | DSHB,<br>H4B4 | Mouse | 1:500 | - |
| GOLPH3 | Abcam<br># ab98023 | Rabbit | - | 1:500 |
| ARFGAP1 | ABclonal<br>#A7118 | Rabbit | 1:1000 | 1:500 |
| IRDye 680 anti-Mouse | LiCOR/926-68170 | Goat | 1:40000 | - |
| IRDye 800 anti-Rabbit | LiCOR/926-32211 | Goat | 1:40000 | - |
| IRDye 800 anti-Goat | LiCOR/926-32214 | Donkey | 1:40000 | - |
| Alexa Fluor 647 anti-Rabbit | Jackson Immuno Research/711-605-152 | Donkey | 1:500 | 1:1000 |
| Alexa Fluor 647 anti-Mouse | Jackson Immuno Research/715-605-151 | Donkey | 1:500 | 1:1000 |
| Alexa Fluor 647 anti-Goat | Jackson Immuno Research/ 705-605-147 | Donkey | - | 1:1000 |
| DyLight 647 anti-Sheep | Jackson Immuno Research/713-605-147 | Donkey | - | 1:1000 |
| anti-Rabbit Cy3 | Jackson Immuno Research/711-165-152 | Donkey | - | 1:1000 |
| anti-Mouse Cy3 | Jackson Immuno Research/715-165-151 | Donkey | - | 1:1000 |
| Alexa Fluor 488 anti-Rabbit | Jackson Immuno Research/711-545-152 | Donkey | - | 1:1000 |
| Alexa Fluor 488 anti-Mouse | Jackson Immuno Research/715-545-151 | Donkey | - | 1:1000 |

### Airyscan Microscopy

Cells were plated on round glass coverslips (12 mm diameter), grown to 80% confluence, washed with PBS and fixed with 4% paraformaldehyde (PFA) (freshly made from 16% stock solution) in PBS for 15 min at RT. Cells were permeabilized with 0.1% Triton X-100 for 1 min, blocked with 50 mM ammonium chloride for 5 min, washed with PBS, and incubated 2 times 10 min each in 1% BSA, 0.1% saponin in PBS. After that, cells were incubated with primary antibodies (diluted in 1% cold fish gelatin, 0.1% saponin in PBS) for 40 min, washed, and incubated with fluorescently conjugated secondary antibodies for 30 min. All the primary and secondary antibodies are listed in Table 1. Hoechst was used to stain chromosomal DNA. Cells were washed four times with PBS, then coverslips were dipped in PBS and water 10 times each and mounted on glass microscope slides using Prolong Gold antifade reagent (Life Technologies). Samples were imaged using 63× oil 1.4 numerical aperture objective with a LSM880 Airyscan Zeiss Laser inverted microscope. Quantitative analysis was performed using ZEN software or ImageJ on single-slice confocal images.

### Transmission Electron Microscopy

The samples were treated according to Valdivia’s protocol [30] with some modifications [31]. Briefly, RPE1 WT, VPS53KO, and VPS54KO cells were plated on a 6 well plate and once the cell reached 90% confluence, 1X fixative was added to equal volume of growing media and incubated at room temperature for 5 min. Next, 0.5X fixative was replaced with 1X fixative (composed of 4% Paraformaldehyde (EMS) and 1% Glutaraldehyde (GA) (EMS) in 0.1 M phosphate buffer of pH 7.4) and incubated for 15 min at RT. The cells were fixed with 2.5% GA, 0.05% malachite green (EMS) in 0.1 M sodium cacodylate buffer of pH 6.8. Cells were washed 4 times 5 min each with 0.1 M sodium cacodylate buffer and post-fixed with 0.5% osmium tetroxide and 0.8% potassium ferricyanide in 0.1 M sodium cacodylate buffer for 40 min at room temperature. The cells were washed again 4 times 5 min each. Then, the cells were incubated with 1% tannic acid for 20 min on ice, washed with buffer and then with water followed by incubation with 0.5% uranyl acetate (UA) at RT for 1 h. After washing with water, cells were scrapped off the plate and transferred to the tubes. 25% to 100% gradual alcohol dehydration was done on ice. Cells were incubated in 100% alcohol 3 times 10 min each, in Propylene oxide (EMS) 3 times 10 min each, and in 1:1 mixture of Propylene oxide and Araldite 502/Embed 812 for overnight. Finally, samples were embedded in Araldite 502/Embed 812 resins (EMS) and hardened at 60°C for 48 h. Ultrathin sections were contrasted by Uranyl Acetate and Reynolds Lead Citrate stains and imaged at 80 kV on FEI Technai G2 TF20 transmission electron microscope. Digital images were acquired with FEI Eagle 4kX USB Digital Camera.

### Quantification of Golgi area

To analyze the *trans*- and *cis*-Golgi area, cells were stained for Rab6 and GM130 and imaged using Airyscan microscopy. Then ImageJ software was used to split the color channels on the individual confocal slices and create binaries. Next, under the function of “Analyze”, the measurement was set to “Area”. The scale of measurement was set for the Golgi particles with size of “2-Infinity” (microns^2) under the “Analyze particles” function. The Golgi “outlines” in both KO cells and control cells were noted and the box-plot graph was made using GraphPad Prism 9.3.0. At least 30 cells were used for quantification of Golgi area per group.

To determine the distance between the *trans*-Golgi and *cis*-Golgi in nocodazole-treated ministacks, a line was drawn in from the center of *cis*-Golgi to the center of *trans*-Golgi within each ministacks in Zen Blue Software. The distance between *cis* and *trans*-Golgi compartments was recorded (n=30).

### Colocalization analysis

Pearson’s correlation coefficient was calculated using “Colocalization” module of Zen Blue software. The colocalization between different proteins was recorded and the graph was made using GraphPad Prism 9.3.0. At least 30 cells were used for quantification of Golgi area per group and Pearson’s correlation coefficient was measured.

### Statistical analysis

All the results are based on at least three different experiments. WB images are representative from three repeats. WBs were quantified using the LI-COR Image Studio software. The error bars for all graphs denote SD. At least 30 cells were used for statistical analysis of Airyscan microscopy. Statistical analysis was done using one-way ANOVA, unpaired t test in GraphPad Prism software.

## Results

### Depletion of GARP complex alters Golgi morphology

Upon microscopic analysis of GARP-deficient RPE1 cells [23] we noticed that Golgi structures in mutant cells looked enlarged and morphologically different from wild type cells. To test if GARP complex subunits knock out (KO) cells have alteration in Golgi size, we stained them using antibodies to *cis*-Golgi protein GM130 and *trans*-Golgi marker Rab6 and examined the size of Golgi compartments using Airyscan superresolution microscopy **(Figure 1A)**. Both *trans*- and *cis*-Golgi areas were significantly increased in VPS54KO cells, while the Golgi increase in VPS53KO cells was less dramatic **(Figure 1B-C)**. To further investigate Golgi changes in GARP-KO cells, we treated them with Nocodazole to disperse Golgi ribbon into ministacks [33, 34]. This approach allowed us to obtain more accurate information about the “Golgi thickness” - the distance between *cis* and *trans*-Golgi compartments **(Figure 1D)**. As expected, Rab6 stained mini-Golgi appeared larger in KO cells and Golgi thickness was significantly increased in both VPS54 and VPS53 deficient cells **(Figure 1E)**. To complement the results obtained with Airyscan microscopy we analyzed GARP-KO cells by Transmission Electron Microscopy (TEM). TEM analysis revealed a tight Golgi ribbon in wild-type (WT) cells whereas, in VPS53KO and VPS54KO cells, the *trans*-Golgi side was severely swollen and fragmented **(Figure 1F-G)**. This indicates that GARP dysfunction in RPE1 cells resulted in an enlarged Golgi structure with severely swollen and fragmented *trans*-Golgi compartments.

**Figure 1.**
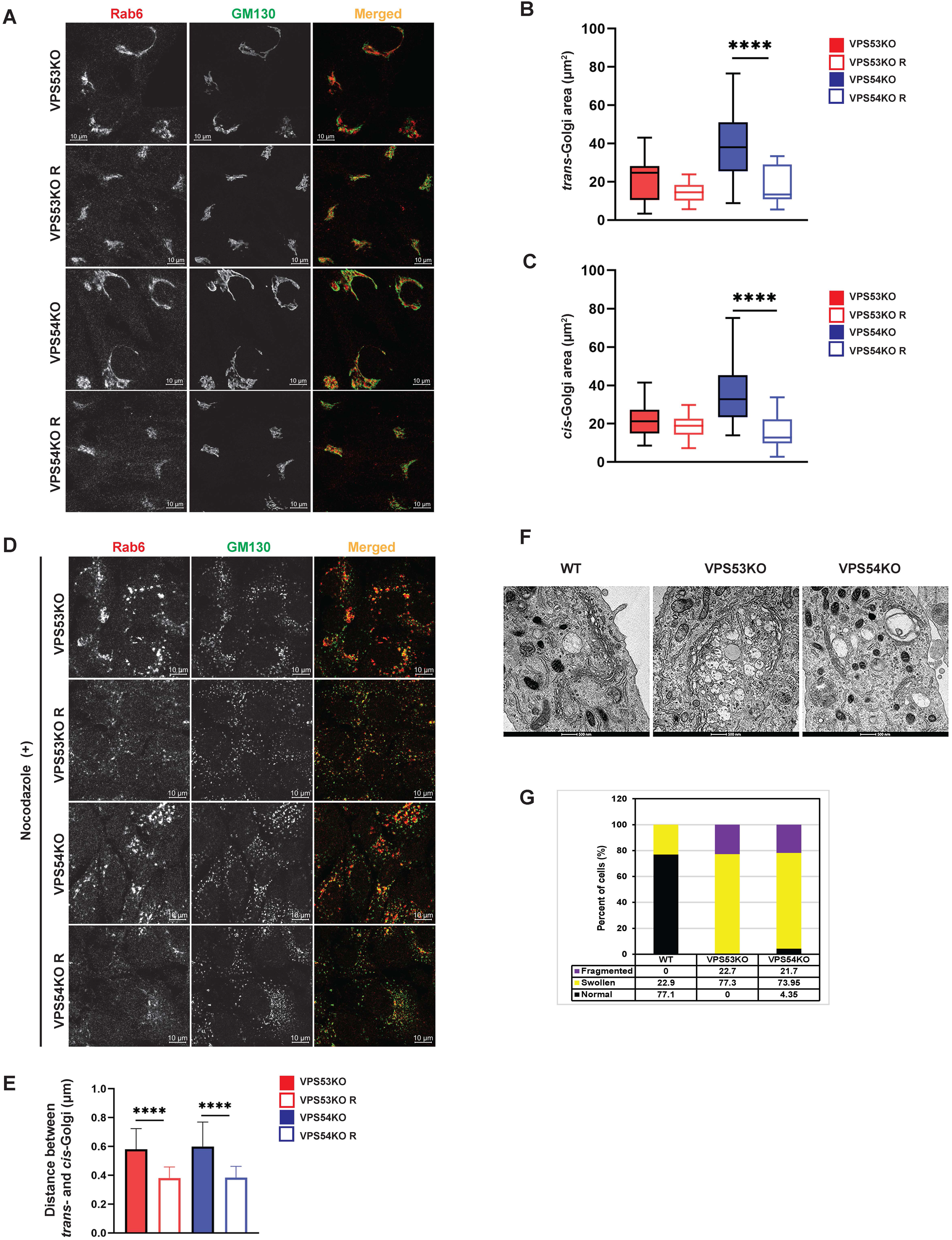
GARP deficiency affects Golgi structure. **(A)** Airyscan microscopy of VPS53KO, VPS53KO R, VPS54KO, and VPS54KO R cells stained for *trans*-Golgi Rab6 and *cis*-Golgi GM130. Individual channels are presented as black and white images, while overlays are presented as RGB images. **(B)** Quantification of *trans*-Golgi area. **(C)** Quantification of *cis*-Golgi area. n=30 cells per group. **(D)** Airyscan microscopy of Nocodazole treated cells, stained for Rab6 and GM130. **(E)** Quantification of the distance between the *cis* (GM130) and *trans* (Rab6) Golgi compartments. **(F)** Transmission electron microscopy of WT, VPS53KO, and VPS54KO RPE1 cells. **(G)** Quantification of normal and abnormal (swollen or fragmented) Golgi in WT, VPS53KO and VPS54KO cells. n=30 cells were imaged per group for the quantification. Statistical significance was calculated using one-way ANOVA in GraphPad prism. \*\*\*\**P* ≤ 0.0001.

### Proteomic analysis revealed depletion of a subset of Golgi proteins in GARP-KO cells

Altered Golgi morphology in GARP-KO cells as well as our previous finding that GARP-KO cells are deficient in several components of Golgi glycosylation machinery [23], triggered the obvious question - how many Golgi proteins depend on the GARP complex for their localization and stability? To answer this question, we utilized label-free protein mass spectrometry proteomic analysis (MS) and compared the abundance of Golgi resident proteins in VPS53KO, VPS54KO, and control cells **(Figure 2A)**. For Golgi isolation, WT and mutant cells were mechanically lysed and fractionated by differential centrifugation to obtain Golgi-enriched 30K membrane pellet. Golgi membranes were further purified from cytoplasmic proteins by floating in the 20-35% Nycodenz density gradient. After centrifugation, gradient fractions were collected for WB analysis. Golgi resident protein B4GALT1 was used as a marker for the Golgi membranes and the majority of B4GALT1 was found in fractions 1-3 both in VPS53KO R cells **(Figure 2B)** and all other WT and COG-deficient cells (A.K. & V.L. unpublished data). Top fractions were also enriched for Golgi proteins STX5 and TMEM165, while the mitochondrial marker COX4 was found in fractions 3-6 (A.K. & V.L. unpublished data). Fractions enriched for B4GALT1 were used for quantitative MS analysis. The MS analysis revealed a significant depletion of 335 proteins in VPS53KO cells, whereas there were 300 proteins depleted in VPS54KO cells **(Figure 2C-D)**. There were 99 common proteins that were depleted in both VPS53KO and VPS54KO cells **(Figure 2D)**. Of these 99 proteins, 22 were Golgi proteins **(Figure 2E**) represented by glycosylation enzymes (orange label), fusion machineries (green), and other Golgi resident proteins (purple). Our previous work [23] demonstrated that two of these proteins, B4GALT1 and TGN46 are severely depleted in GARP-KO cells. While most GARP-sensitive proteins localize in *trans*-Golgi compartments, some of them, like GLG1/MG-160, GALNT1, MAN1A2 [35, 36], are reported to localize in *cis/medial* Golgi cisternae, indicating that GARP dysfunction is affecting the entire Golgi complex.

**Figure 2.**
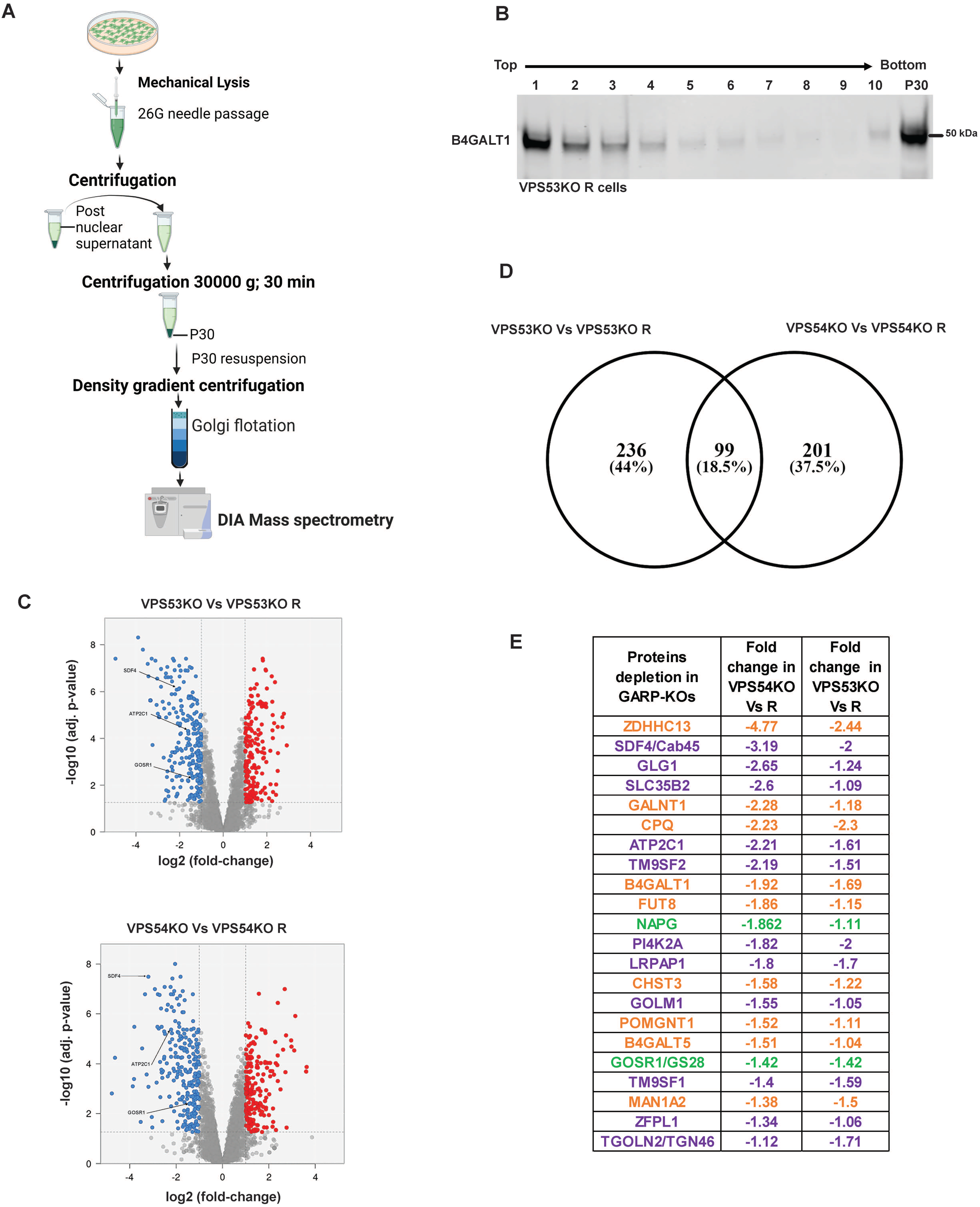
Quantitative Mass spectrometry (MS) analysis of Golgi-enriched membranes isolated from GARP-KOs revealed depletion of multiple Golgi proteins. **(A)** Schematic of the major steps during preparation of Golgi samples for MS analysis. **(B)** WB for Golgi marker B4GALT1 in membranes isolated from VPS53KO R cells and separated by the Nycodenz flotation gradient. **(C)** Volcano plot analysis of relative protein abundance in VPS53KO and VPS53KO R (Top) and VPS54KO and its VPS54KO R cells (Bottom). Blue circles represent the proteins significantly depleted in GARP-KO cells. Red circles represent proteins significantly increased in GARP-KO cells. Gray circles are proteins with no significant differences. **(D)** Venn diagram demonstrating common proteins depleted in both VPS53KO and VPS54KO cells. **(E)** List of Golgi proteins significantly depleted in both VPS53KO and VPS54KO cells. Proteins with color orange are Golgi enzymes, purple are Golgi resident proteins, and Green are Golgi SNAREs.

### GARP dysfunction affects Golgi calcium pump and calcium binding protein SDF4

To validate MS results we tested the localization and abundance of several Golgi proteins, first focusing on two key players in Golgi Ca^2+^ homeostasis, SDF4/Cab45 and ATP2C1/SPCA1. SDF4 is a *trans*-Golgi network luminal calcium-binding protein that promotes sorting of a subset of secretory proteins at the *trans*-Golgi Network (TGN) [37]. To verify the SDF4 MS results, we co-stained VPS53KO, VPS54KO and control cells with SDF4 and a GARP-independent *trans*-Golgi protein Golgin97/GOLGA1 [38] **(Figure 3A)**. Airyscan microscopy revealed a significant decrease in both SDF4 Golgi signal intensity and colocalization of SDF4 with Golgin97 supporting MS results **(Figure 3B)**. Furthermore, the total SDF4 protein level was significantly decreased in both VPS53KO and VPS54KO cells (**Figure 3C-D)**. Ca^2+^/Mn^2+^ ATPase ATP2C1/SPCA1 [39, 40] is another Golgi protein that was depleted in the Golgi membranes isolated from GARP-KOs. Like in the case of SDF4, we found a significant reduction in the total protein abundance of ATP2C1 in GARP-KOs **(Figure 3E-F)**. In summary, our analysis revealed two proteins related to Ca^2+^ Golgi homeostasis are depleted in GARP-KO cells.

**Figure 3.**
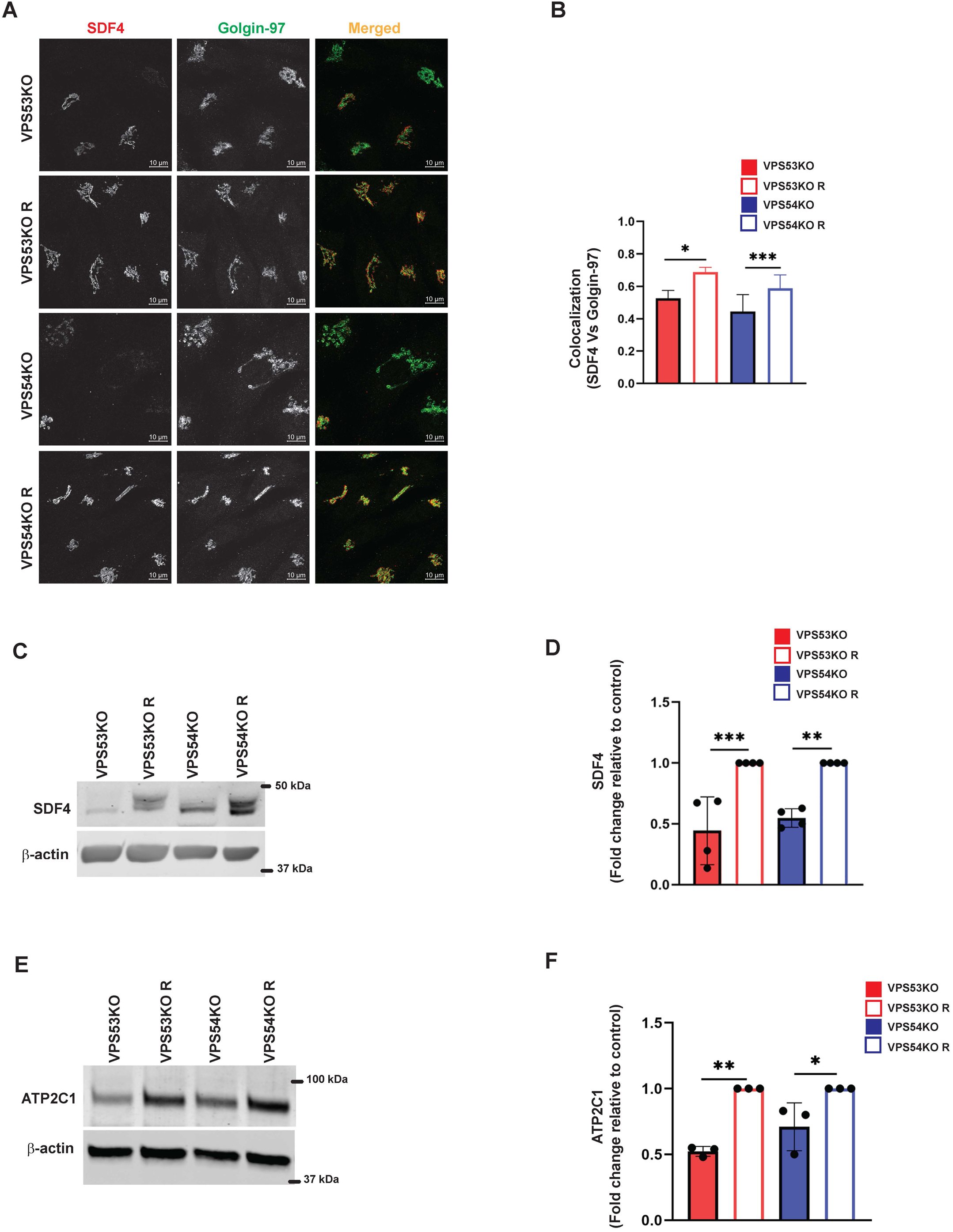
GARP dysfunction alters the Ca2+ binding SDF4/Cab-45 and Ca^2+^-transporting ATP2C1/SPCA1 Golgi proteins. **(A)** Airyscan microscopy of VPS53KO, VPS53KO R, VPS54KO and VPS54KO R cells co-stained with SDF4 and Golgin97. **(B)** The graph shows the quantification of SDF4 and Golgin97 colocalization. Values in the bar graph represent the mean ± SD from the co-localization between SDF4 and Golgin97 from 50 different cells. **(C)** WB analysis of SDF4 in GARP-KO and rescued cells. **(D)** Quantification of SDF4 WBs from three independent experiments. **(E)** WB analysis of ATP2C1 in GARP-KO and rescued cells. **(F)** Quantification of ATP2C1 blots from three independent experiments. β-actin was used as the internal loading control. Values in bar graph represent the mean ± SD from at least three independent experiments. Statistical significance was calculated using one-way ANOVA. \*\*\**P* ≤ 0.001, \*\**P* ≤ 0.01, \**P* ≤ 0.05.

### GARP depletion alters expression and localization of intra-Golgi v-SNAREs

Another key component of Golgi homeostasis significantly depleted in the GARP-deficient Golgi was Qb-SNARE GOSR1/GS28. We validated the MS results both by IF **(Figure 4A-B)** and WB analysis **(Figure 4C-D)**. GOSR1 was mostly localized in the Golgi in control cells and was displaced from GM130-defined Golgi region in VPS53KO and VPS54KO cells confirming that the intracellular localization of this Golgi SNARE is severely altered in GARP-KOs. We next tested if the change in Golgi localization of GOSR1 affects its total protein level. Indeed, we observed a significant depletion in the abundance of GOSR1 in total cell lysates of GARP-KOs **(Figure 4C-D)**. GOSR1 works in the STX5/GOSR1/BET1L/YKT6 SNARE complex to facilitate fusion of intra-Golgi recycling vesicles [41]. We have previously shown that Qc-SNARE, BET1L/GS15, is depleted in GARP-KO cells [42]. Similar to GOSR1 results, severe mislocalization of BET1L was observed in both VPS54KO and VPS53KO cells **(Figure 4E-F)**. Proteomics data indicated that two other SNAREs of this complex, YKT6 and STX5, are not depleted in the Golgi-enriched membranes from GARP mutants, indicating that GARP deficiency only affects putative v-SNARE proteins. This is in line with the data obtained in COG-deficient cells [43]. GOSR1 and BET1L are two major Golgi Q-SNAREs involved in the intra-Golgi retrograde trafficking of Golgi resident proteins [41] and their mislocalization and degradation could be responsible for Golgi defects in GARP-KO cells. To test this possibility, we created RPE1 cells deleted for GOSR1 and BET1L **(Figure 5A)**. Deletion of these Golgi SNAREs did not alter RPE1 cell growth (data not shown), similar to SNARE KO results in HEK293T cells [44]. To test if SNARE depletion would compromise the stability of Golgi resident proteins, the total protein abundance of SDF4, B4GALT1 and TGN46 was compared between WT, SNAREs-KO (BET1LKO, GOSR1KO) and GARP-KO (VPS54KO) cells **(Figure 5B-C)**. Although the abundance of SDF4 **(Figure 5B, C**), B4GALT1 **(Figure 5D, E)** and TGN46/TGOLN2 **(Figure 5F, G)** was reduced in SNARE-KO cells, GARP depletion demonstrated a significantly greater decrease in the abundance on all three Golgi resident proteins, indicating that depletion of Golgi SNAREs alone could not explain Golgi defects in GARP-KO cells **(Figure 5B-G)**. Another important phenotype observed in GARP-KO cells is their alteration in Golgi size as shown in **Figure 1A**. We, therefore, examined Golgi morphology by staining the WT, BET1LKO, GOSR1KO and VPS54KO cells with *trans*-Golgi marker Rab6 and *cis*-Golgi marker GM130 **(Figure 5H)**. Airyscan microscopy analysis revealed that the Golgi apparatus was more enlarged in VPS54KO cells compared to SNARE-KO cells confirming that GARP-KO Golgi-related defects are more severe than defects observed in SNAREs-KOs.

**Figure 4.**
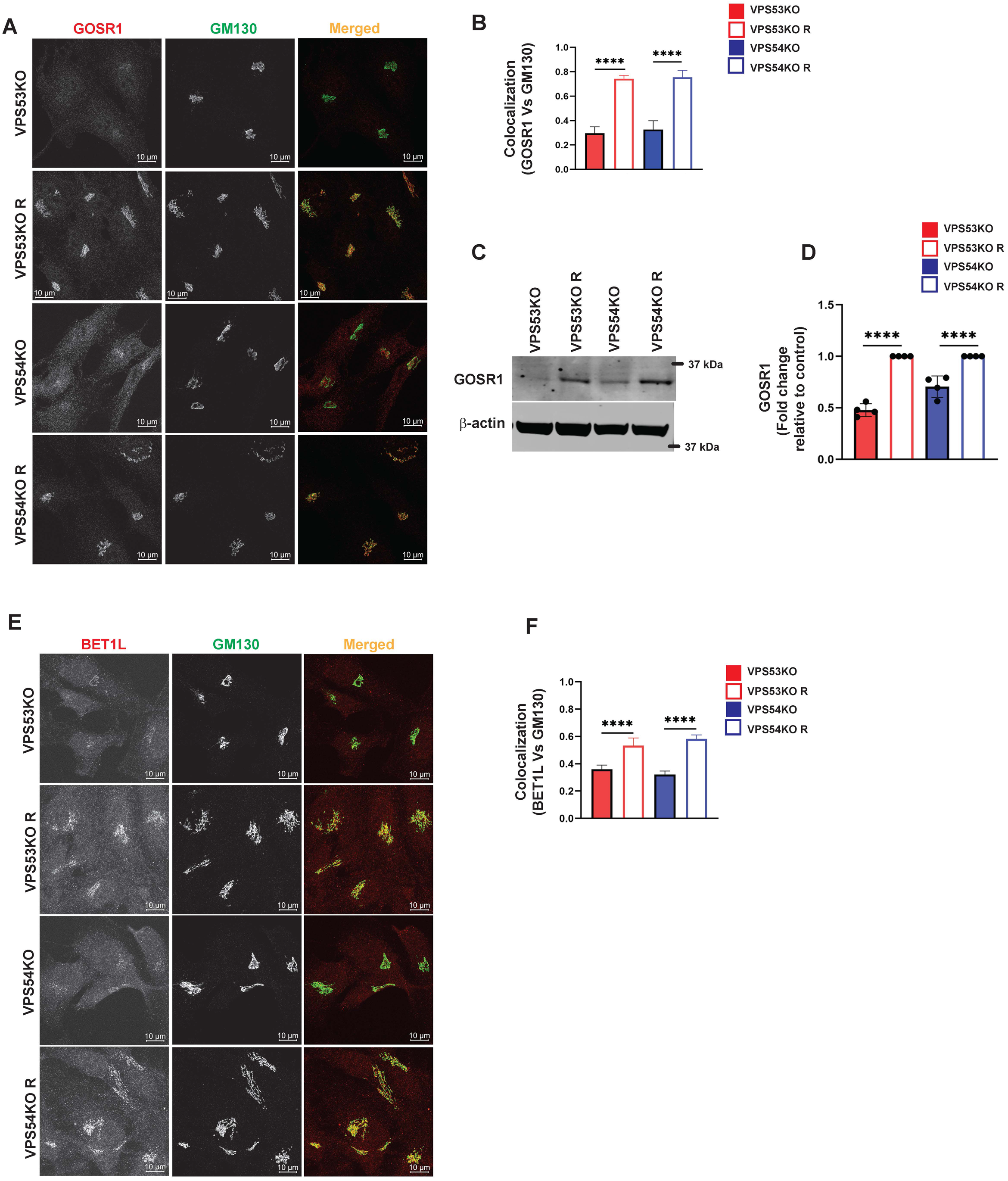
GARP-KO alters localization and expression of intra-Golgi v-SNAREs GOSR1 and BET1L. **(A)** Airyscan microscopy of GARP-KOs and control cells co-stained for GOSR1 and GM130. **(B)** The graph shows the quantification of GOSR1 and GM130 colocalization. Values in the bar graph represent the mean ± SD from the colocalization between GOSR1 and GM130 from 40 different cells. **(C)** WB analysis of GOSR1 protein level in VPS53KO, VPS54KO, and their controls. β-actin was used as internal loading control. **(D)** Quantification of the blots from four independent experiments. **(E)** Airyscan imaging of VPS53KO, VPS53KO R, VPS54KO, VPS54KO R cells co-stained for BET1L and GM130. **(F)** Colocalization analysis between BET1L and GM130 using Pearson’s correlation coefficient. n= 40 cells were imaged per group for the quantification. Statistical significance was calculated using one-way ANOVA. \*\*\*\**P* ≤ 0.0001.

**Figure 5.**
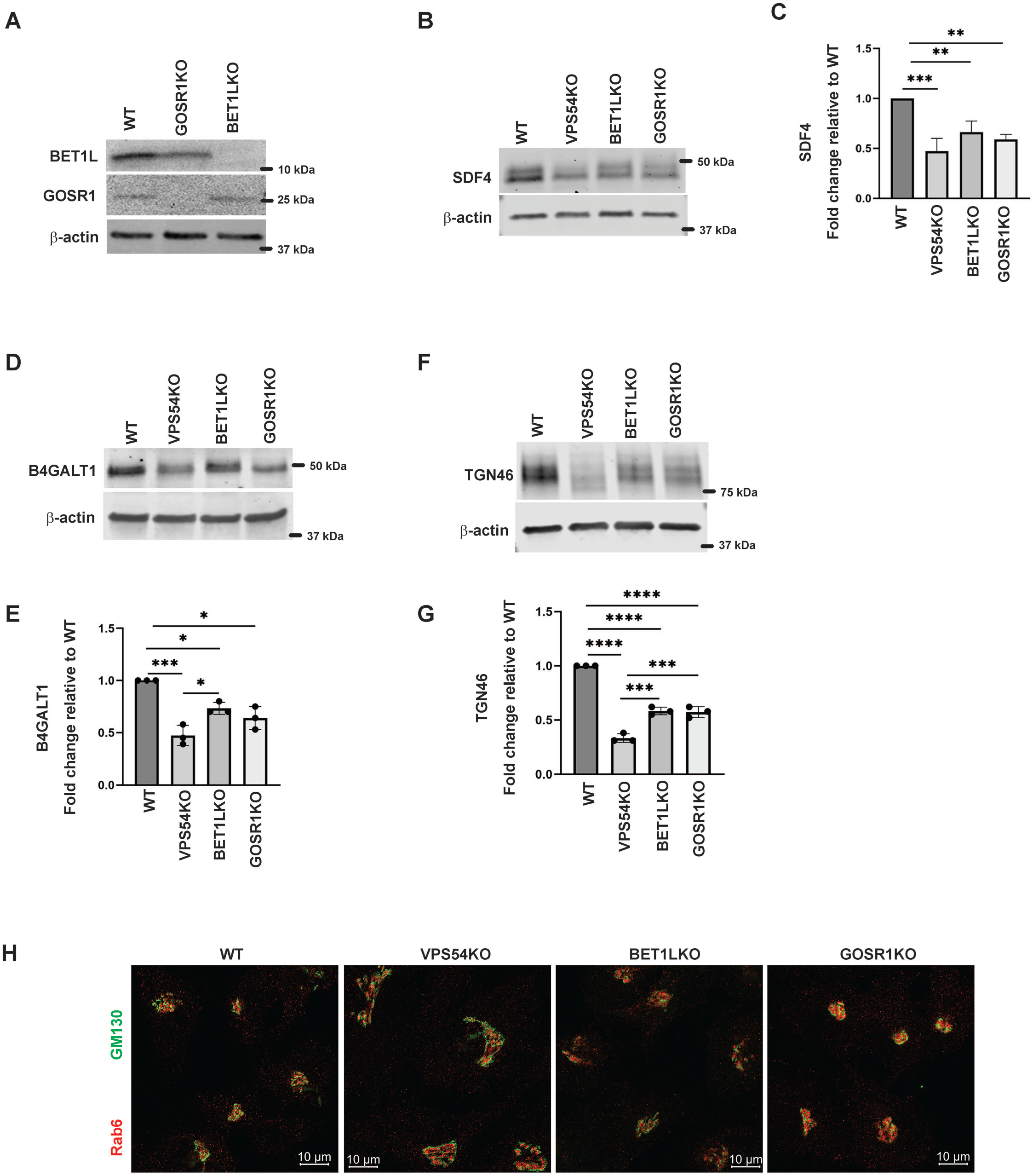
GARP-KO is more detrimental to Golgi residents than deletion of Golgi v-SNAREs. **(A)** Testing of GOSR1KO and BET1LKO by WB analysis. **(B)** Total cellular lysates were prepared from RPE1 WT, VPS54KO, BET1LKO, and GOSR1KO cells and total protein abundance was analyzed by WB for SDF4, **(D)** B4GALT1, and **(F)** TGN46. Normalization was performed using β-actin. Quantification of relative total protein level of **(C)** SDF4, **(E)** B4GALT1, and **(G)** TGN46. Statistical analysis was done from three independent blots, where \*\*\*\**P* ≤ 0.0001, ***P ≤ 0.001, \*\**P* ≤ 0.01, *P ≤ 0.05. **(H)** Airyscan microscopy of RPE1 WT, VPS54KO, BET1LKO, and GOSR1KO cells stained for Rab6 and GM130.

### COPI vesicle coat proteins relocate to ERGIC in GARP-KO cells

Severe depletion of intra-Golgi trafficking SNAREs GOSR1 and BET1L in GARP-KO cells should lead to defects in a vesicle fusion step but TEM analysis of GARP-KO cells did not reveal any significant accumulation of vesicular structures **(Figure 1F)**, suggesting that formation of transport vesicles in mutant cells could be altered as well. To test this possibility, we analyzed the localization of vesicle coat machinery in GARP-KO cells. COPI complex consists of seven subunits - beta, gamma, delta, zeta, alpha, beta prime and epsilon [45]. We performed IF and investigated both proximal membrane coat proteins (COPB1, COPG1) as well as distal membrane coat subunit COPB2. We observed a significant reduction in juxtanuclear localization of both layers of COPI coat **(Figure 6A-D)** in GARP-KOs, with the majority of COPI signal appearing as peripheral dots. To determine if COPI is relocalized to pre- or post-Golgi compartments we co-stained cells with COPI subunits and ERGIC-53 and found their colocalization increased in VPS54KO cells **(Figure 6 E-G)**. Co-staining with COPG1 and SNX1 did not reveal any significant colocalization between COPI and endosomes (A.K. unpublished data).

**Figure 6.**
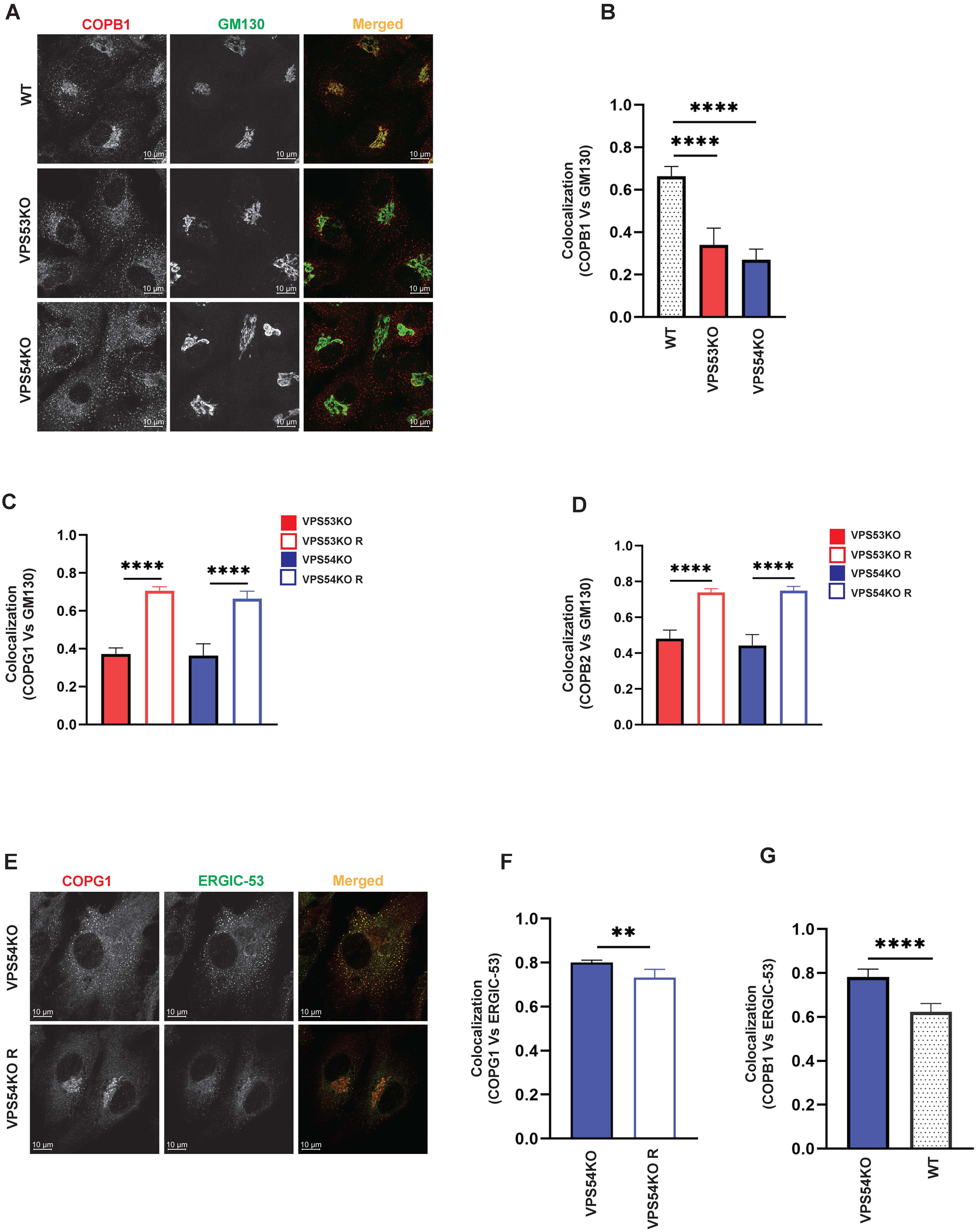
COPI subunits are displaced from the Golgi in GARP-KO cells. **(A)** Airyscan microscopy of GARP-KO cells and control cells stained with antibody to coatomer subunit COPB1. Colocalization analysis of COPB1 and GM130 **(B)**, COPG1 and GM130 **(C)**, COPB2 and GM130 **(D)** using Pearson’s correlation coefficient. n=40 cells used for colocalization analysis per group. **(E)** Airyscan microscopy of COPG1 and ERGIC-53 in VPS54KO and VPS54KO R cells. **(F)** Colocalization analysis of COPG1 and ERGIC-53 and **(G)** COPB1 and ERGIC-53 using Pearson’s correlation coefficient. n=40 cells used for colocalization analysis. Statistical significance was calculated using one-way ANOVA. \*\*\*\**P* ≤ 0.0001, \*\**P* ≤ 0.01.

### GARP dysfunction leads to mislocalization of COPI accessory proteins

Our previous study [23] and the current proteomics data revealed depletion of several Golgi glycosyltransferases (B4GALT1, GALNT1, ST6GAL1 and MGAT1) in GARP-KO cells. GOLPH3 plays a crucial role in retrograde intra-Golgi trafficking of glycosyltransferases [46]. It is shown to bind the cytoplasmic tails of Golgi enzymes and packages them into recycling COPI vesicles [47]. ARFGAP1 promotes the formation of COPI vesicles [48, 49]. Both GOLPH3 and ARFGAP1 are peripheral membrane proteins that were stripped from membranes during flotation in the Nycodenz gradient and not detected by MS analysis in the Golgi-enriched membranes. But since the localization of COPI was found to be GARP-sensitive, the localization of GOLPH3 and ARFGAP1 in GARP deficient cells was tested. We found a decrease in Golgi localization of both GOLPH3 **(Figure 7A-B)** and ARFGAPs **(Figure 7C-D)**, indicating that COPI accessory proteins also require GARP activity for their proper localization. However, unlike intra-Golgi v-SNAREs, the total cellular level of ARFGAP1 protein was increased **(Figure 7E-F)**, while GOLPH3 expression was not altered (A.K. unpublished data) in GARP-KO cells.

**Figure 7.**
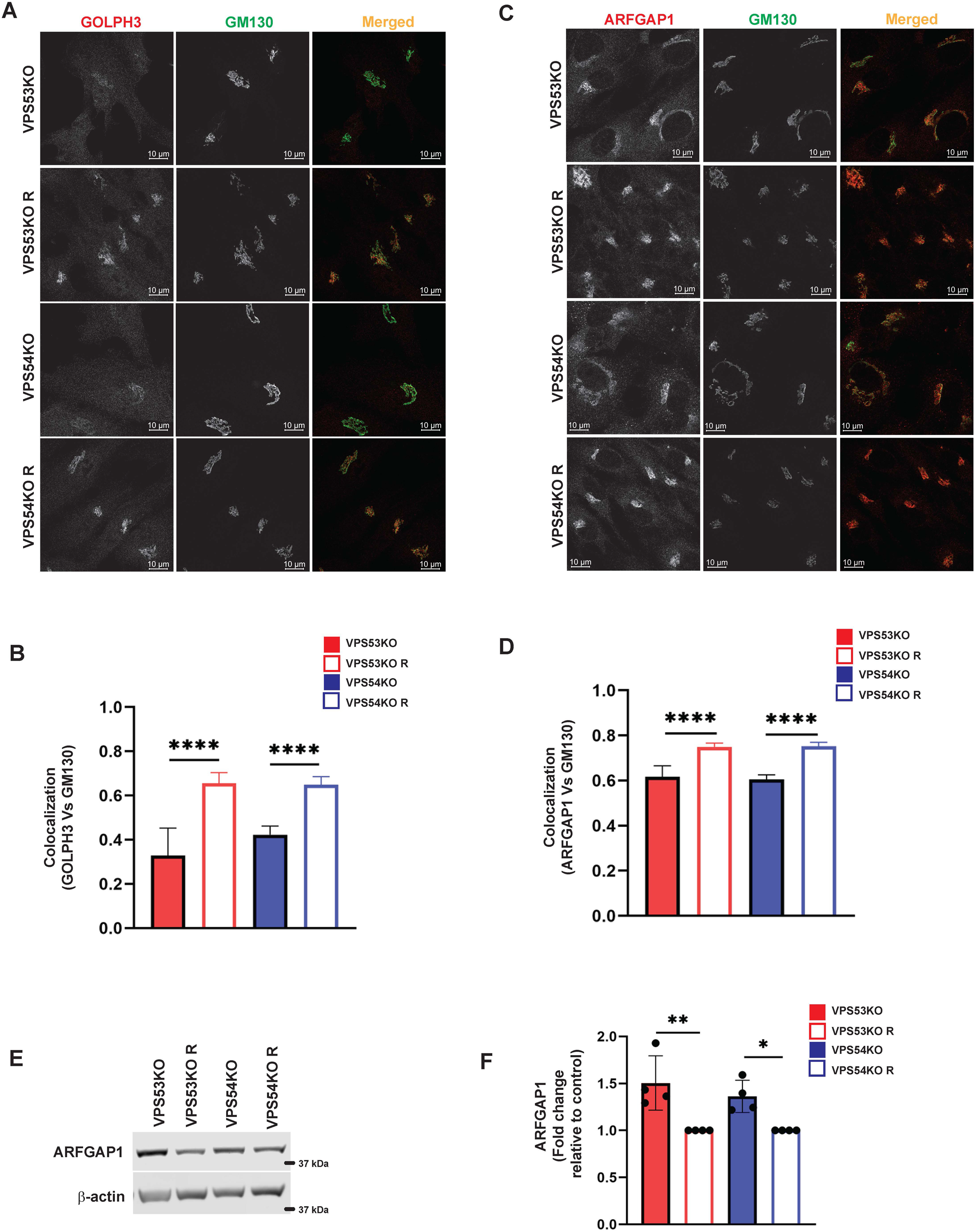
GARP dysfunction resulted in mislocalization of COPI adapter protein GOLPH3 and ARFGAP1. **(A)** Airyscan microscopy of VPS53KO, VPS53KO R, VPS54KO, VPS54KO R cells stained with GOLPH3 and GM130. **(B)** Colocalization analysis between GOLPH3 and GM130 using Pearson’s correlation coefficient. n=40 cells were used for colocalization analysis. **(C)** VPS53KO, VPS53KO R, VPS54KO, and VPS54KO R cells were stained for ARFGAP1 and GM130. **(D)** Colocalization analysis of ARFGAP1 and GM130 using Pearson’s correlation coefficient. n= 40 cells were used for colocalization analysis. **(E)** WB analysis of ARFGAP1 total protein abundance in GARP-KO and control cells. **(F)** Quantification of total protein abundance of ARFGAP1 in GARP-KOs and control cells. β-actin was used as internal loading control. \*\*\*\**P* ≤ 0.0001, \*\**P* ≤ 0.01, \**P* ≤ 0.05.

### Localization of ARFGEFs is severely affected in GARP-KO cells

COPI binding to Golgi membrane requires activation of ARF GTPases, which is facilitated by ARFGEF proteins [50]. GBF1 is an ARFGEF found in *cis*-Golgi and ERGIC, [51]. To test if GARP-KOs have any effect in ARFGEF that could result in prevention of COPI coats assembly, we stained cells with GBF1 and *cis*-Golgi marker GM130 **(Figure 8A)**. Interestingly, we found a significant displacement of GBF1 from the *cis*-Golgi in GARP-KO cells. GBF1 displacement was somewhat heterogeneous with larger cells displaying a more severe phenotype. GBF1 displacement from the Golgi was further confirmed by colocalization analysis between GBF1 and GM130 using Pearson’s correlation coefficient **(Figure 8B)**. This phenotype was not cell-line dependent as both HeLa **(Figure 8C)** and HEK293T **(Figure 8D)** GARP-depleted cells showed decrease in colocalization of GBF1 with GM130. We not only found GBF1 was depleted in GARP-KOs, but we also found BIG1, an ARFGEF known to function in *trans*-Golgi and endosomes [52] was altered **(Figure 8E-F)**. In VPS54KO cells, we observed that BIG1 was mostly relocated to bright peripheral dots **(Figure 8E)**, and we were inquisitive about the nature of that compartment. Co-staining cells with BIG1 and ERGIC-53 **(Figure 9A-B)** or with BIG1 and SNX1 **(Figure 9C)** did not reveal any significant increase in colocalization, indicating that BIG1 did not relocalize to the ERGIC or sorting endosomes. Furthermore, we tested if BIG1 is localized to LAMP2 compartment in GARP-KOs **(Figure 9D)**. Surprisingly, we found a significant increase in colocalization of BIG1 to LAMP2 compartment in GARP-KOs **(Figure 9E)**. Similar results were obtained by co-staining cells with BIG1 and another marker of endolysosomal compartment, CD63 **(Figure 9F)**. Again, a significant increase in colocalization between CD63 and BIG1 was observed **(Figure 9G)**. Taken together, these results indicated that GARP depletion affects Golgi localization of ARFGEF GBF1 and BIG1, and BIG1 is displaced to endolysosomal compartment.

**Figure 8.**
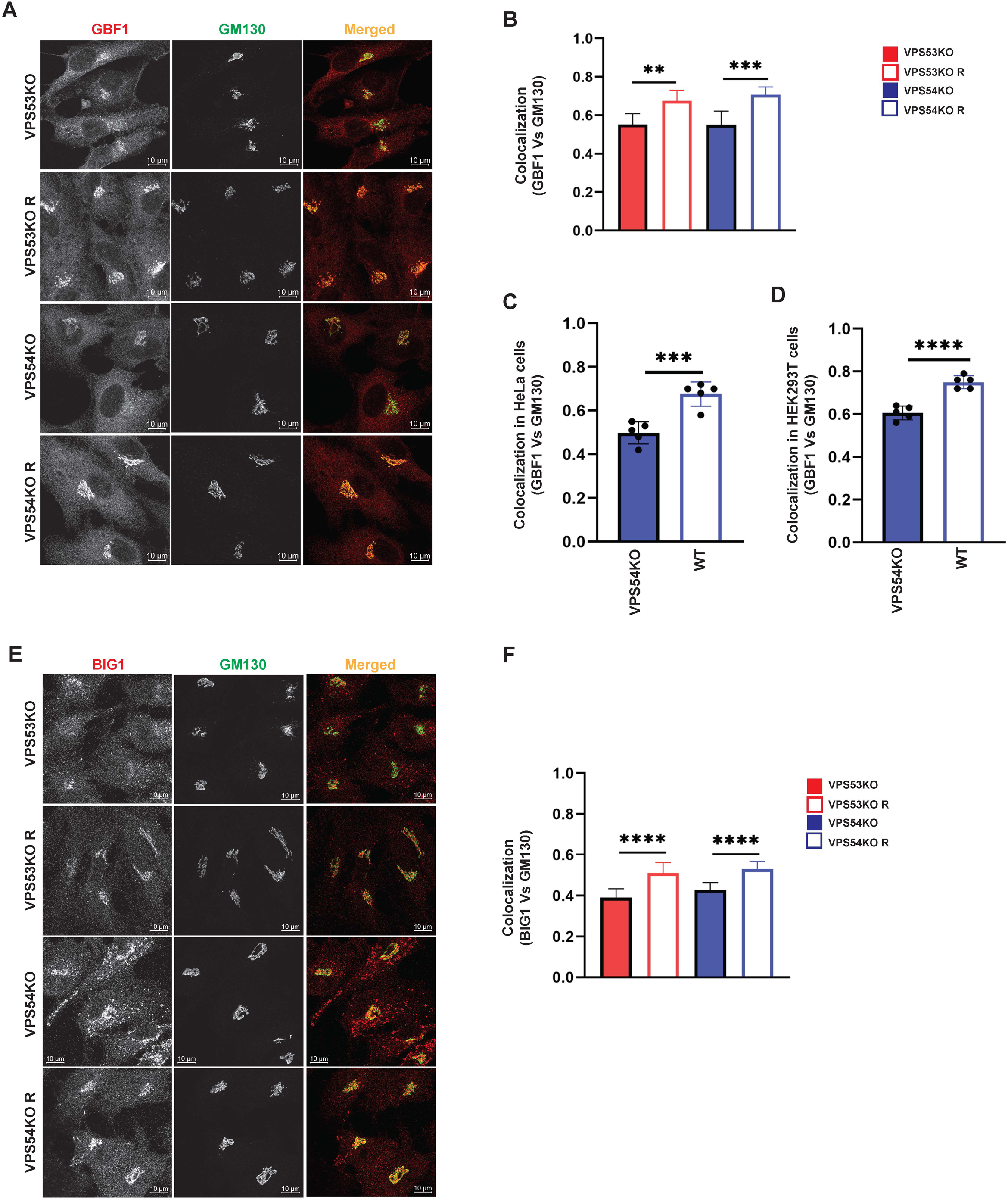
ARFGEFs GBF1 and BIG1 are displaced to off-Golgi compartments in GARP-KO cells. **(A)** Airyscan microscopy of GARP-KO and control cells co-stained with antibodies to GBF1 and GM130 in RPE1 cells. **(B)** Colocalization analysis of GBF1 and GM130 in RPE1 cells. **(C)** Colocalization analysis of GBF1 and GM130 in HeLa cells. **(D)** Colocalization analysis of GBF1 and GM130 in HEK293T cells. **(E)** Airyscan microscopy of GARP-KO and control cells co-stained with antibodies to BIG1 and GM130 in RPE1 cells. **(F)** Colocalization analysis of BIG1 and GM130 using Pearson’s correlation coefficient. n=50 cells were used for colocalization analysis. \*\*\*\**P* ≤ 0.0001, \*\*\**P* ≤ 0.001, \*\**P* ≤ 0.01.

**Figure 9.**
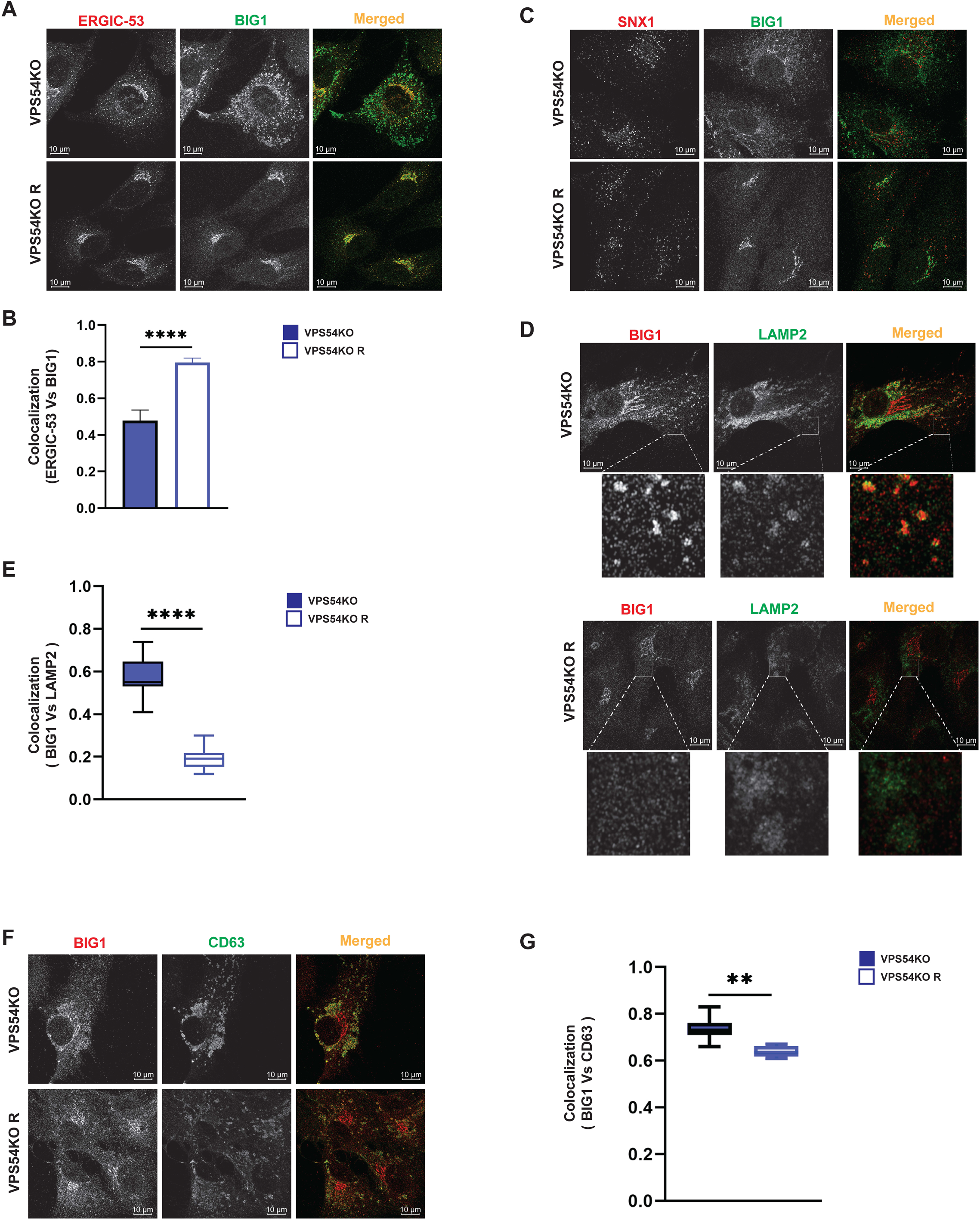
BIG1 is mislocalized to endolysosomal compartment in GARP-KOs. Airyscan microscopy of VPS54KO and VPS54KO R cells stained for BIG1 and ERGIC-53 **(A)**, BIG1 and SNX1 **(C)**, BIG1 and LAMP2 **(D)**, BIG1 and CD63 **(F)**. Colocalization analysis between BIG1 and ERGIC-53 **(B)**, BIG1 and LAMP2 **(E)**, BIG1 and CD63 **(G)** by Pearson’s correlation coefficient analysis. n=30 cells. \*\*\*\**P* ≤ 0.0001, \*\**P* ≤ 0.01.

## Discussion

In this study, we have extended our investigation of the Golgi defects in human cells completely depleted for the GARP complex. The summary of observed GARP-related Golgi defects is presented in **Figure 10**. Microscopy and proteomic approaches revealed severe alterations of the Golgi structure in GARP-KO cells that coincided with a significant depletion of a subset of Golgi resident proteins. Golgi morphological alterations in GARP deficient cells (volume expansion of all subcompartments) were distinct from the severe fragmented Golgi phenotype observed in cells depleted for the other two Golgi vesicle tethering complexes COG [43, 53, 54] and ZW10/Dsl10 [54, 55], indicating that GARP depletion affects a specific set of Golgi proteins. Indeed, Golgi proteomics revealed a distinct set of Golgi resident proteins affected in GARP-KO cells. As predicted from our previous study [42], the set of GARP-sensitive proteins includes several glycosylation enzymes. We also discovered that KO of GARP complex subunits affects the calcium pump ATP2C1 as well as calcium binding protein SDF4. Furthermore, we found that localization of key elements of intra-Golgi trafficking machineries including v-SNAREs, COPI proteins, ARFGEFs are also severely affected in GARP-KO cells.

**Figure 10.**
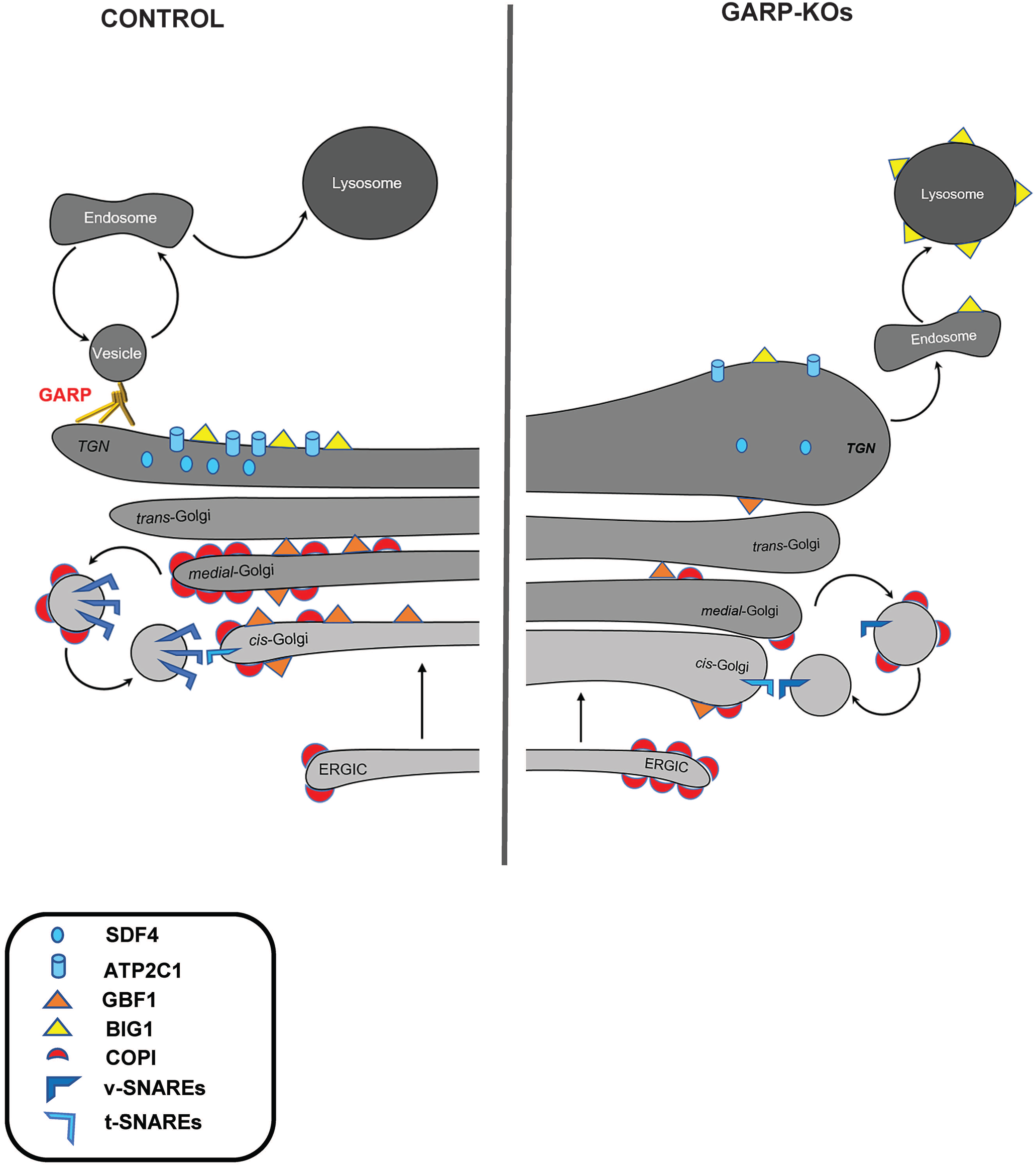
GARP dysfunction alters Golgi homeostasis and Golgi trafficking machineries. A cartoon depicting Golgi trafficking machinery in WT and GARP-KO cells. Proper maintenance of Golgi structure, calcium channel and Golgi trafficking machinery including SNAREs, COPI coats, ARFGEFs in control cells (Left panel). Disruption of the Golgi structure, calcium channel, and Golgi trafficking machineries in GARP-KO cells (Right panel).

SDF4/Cab45 is a calcium-binding luminal Golgi resident protein that is responsible for sorting of specific cargo proteins at the TGN [37, 56-59]. SDF4 depletion in the Golgi can be due to several reasons. First, SDF4 could be cycling between TGN and endosomal compartment in GARP-dependent manner and failed to return to TGN in GARP depleted cells. Second, SDF4 retention in the TGN requires high Ca^2+^ concentration in the Golgi lumen [60], which could be altered in GARP-KO cells. Interestingly, in both scenarios we expected to find an increased secretion of SDF4 in GARP-KO cells. The third possibility is that GARP deficiency is forcing the displacement of SDF4 into other secretory/endolysosomal compartments. Our preliminary results (A.K. unpublished data) indicate a decrease of SDF4 in the secretome from GARP-KO cells which suggests that the decrease in SDF4 cellular levels is likely caused by its missorting and degradation to the lysosomal compartment. SDF4 missorting is likely to relate to the depletion of Golgi calcium pump ATP2C1/SPCA1. ATP2C1 pumps Ca^2+^ into the TGN lumen and defect in ATP2C1 results in missorting of secretory cargo [39, 40]. Mutation of ATP2C1 gene is associated with Hailey-Hailey disease [61, 62]. Depletion of ATP2C1 in GARP-KOs could also be connected to the alteration in TGN morphology. In support of this hypothesis, Micaroni et al. showed that correct ATP2C1 functioning is critical for intra-Golgi trafficking and maintenance of Golgi structure [63-65].

GARP was shown to regulate formation and/or stability of TGN STX16/STX6/VTI1A/VAMP4 SNARE complex [10, 66], but surprisingly we did not find any components of STX16 complex among proteins depleted from the Golgi membranes in GARP-KO cells. Instead, GOSR1, v-SNARE of intra-Golgi STX5/GOSR1/BET1L/YKT6 [67] complex was severely depleted in GARP-KO cells. BET1L, most likely because of its small size, was not detected in the proteomic studies, but our analysis indicated mislocalization of this Golgi v-SNARE in GARP-KO cells. The total BET1L protein level was also decreased [23]. This indicates that GARP could be involved in regulation and/or stability of STX5/GOSR1/BET1L/YKT6 SNARE complex. Interestingly, this SNARE complex has been implicated not only in intra-Golgi [41, 68] but also in the endosome to TGN transport [69]. Therefore, one explanation for the loss of GOSR1 and BET1L in GARP deficient cells is their inability to recycle back to the Golgi from the endosomal compartment. Importantly, the complete knock out of GOSR1 or BET1L is less deleterious to cells as compared to the loss of the GARP complex, indicating that the loss of v-SNARE alone could not explain all Golgi phenotypes observed in GARP deficient cells.

Indeed, we discovered that the entire COPI vesicle budding machinery is significantly mislocalized in GARP deficient cells. The majority of COPI was no longer Golgi localized. Instead, it showed an increased co-localization with the ER-Golgi (ERGIC) intermediate compartment. Two COPI associated proteins, GOLPH3 and ARFGAP1 are also displaced from the Golgi in GARP-KO cells, not to the ERGIC, but likely dissociate to cytoplasm. GOLPH3 plays a crucial role in sorting of a subset of Golgi resident glycosyltransferases back to Golgi by binding to the cytoplasmic tails of Golgi glycosyltransferases to package them into recycling COPI vesicles [46, 47, 70-73], and its displacement from the Golgi in GARP deficient cells could be responsible for some glycosylation defects in mutant cells. Displacement of GOLPH3, however, could not explain the instability of B4GALT1 and MGAT1 in GARP-KO cells [42], since GOLPH3 is not responsible for their Golgi retention. So, our next question was what would be the reason for decrease in Golgi localization of COPI coats?

GBF1 is a cis-Golgi ARF GEF that activates ARF GTPases and takes part in COPI recruitment, Golgi integrity and secretory traffic whereas inactivation or depletion of GBF1 inhibits these processes [74-76]. We were unable to investigate the changes in localization of the endogenous Arf1 in GARP-KO cells, but significant displacement of GBF1 from Golgi membranes is likely to cause Arf1 and COPI depletion from the Golgi membranes. One of the reasons for depletion of GBF1 and GOLPH3 in the Golgi could be an alteration of membrane lipid content in GARP-KO cells. It has been demonstrated previously that inhibitors of PI4P synthesis prevent the recruitment of GBF1 to Golgi membranes [77]. PI4P is also essential for GOLPH3 binding to membranes [78]. Indeed, MS analysis revealed a a significant decrease in Golgi associated PI4K2a kinase, supporting the possibility that PI4P Golgi content is altered in GARP-KO cells. In addition, GARP mutations in yeast and mouse models result in sphingolipid abnormalities [20] [79]. The effect of GARP depletion on the lipid content of the Golgi will be investigated in the future.

Not only GBF1 was depleted from the Golgi in GARP-KOs, we also found BIG1, an ARFGEF is known to function in *trans*-Golgi to endosome [80] was altered. This is in consistent with the finding that inactivation of GBF1 by inserting mutation or treating with GBF1-selective drug golgicide (GCA) inhibited membrane recruitment of BIG1 and BIG2 [81]. We also observed GGA2 recruitment being depleted in GARP-KO RPE1 cells (unpublished data). This is consistent GBF1 is required for GGA recruitment to Golgi membranes and plays a role in the proper processing and sorting of lysosomal cargo [82]. We observed that BIG1 in GARP-KOs is mislocalized to endolysosomal compartments. Although the exact mechanisms of GARP-dependent relocalization of COPI machinery will require additional investigation, we propose that the dysregulation of COPI machinery along with degradation of intra-Golgi v-SNAREs and alteration of Golgi Ca2^+^ homeostasis are the major driving factors for the instability of Golgi resident proteins and glycosylation defects in GARP deficient cells.

## Supporting information

Golgi proteomics data for Figure 2

## Abbreviations used

B4GalT1: Beta-1,4-galactosyltransferase 1
COG: Conserved Oligomeric Golgi
GARP: Golgi-associated retrograde protein
SNARE: SNAP (Soluble NSF Attachment Protein) Receptor
TGN: trans-Golgi Network

## Conflict of Interest

No conflict of interests was declared.

## Author Contributions

A.K. wrote the article and made substantial contributions to conception and design, acquisition of data, analysis and interpretation of data. T.K. edited the article, performed experiments for Figure 2 and interpreted the data. I.P. edited the article, performed experiments for Figure 1 and interpreted the data. Z.D. edited the article, performed experiments for Figure 5 and interpreted the data. V.L. wrote the article and made substantial contributions to conception and design.

## Funding

This work was supported by the National Institute of Health grant R01GM083144 and by UAMS Easy Win Early Victory grant program for Vladimir Lupashin

## Acknowledgments

We are thankful to Juan S. Bonifacino, Rainer Duden, Taroh Kinoshita and others who provided reagents and cell lines. We would also like to thank the UAMS IDeA National Resource for Quantitative Proteomics, sequencing, flow cytometry and digital microscopy core facilities for the use of their equipment and expertise. We are grateful to all members of Lupashin’s lab for comments on the manuscript.

## Notes

### Competing Interest Statement

The authors have declared no competing interest.

